# Global epistasis in budding yeast driven by many natural variants whose effects scale with fitness

**DOI:** 10.1101/2025.05.28.656710

**Authors:** Ilan Goldstein, Joseph J. Hale, Ian M. Ehrenreich

## Abstract

Global epistasis is a phenomenon in which the effects of genetic perturbations depend on the fitness of the individuals in which they occur. In populations with natural genetic variation, global epistasis arises from interactions between perturbations and polymorphic loci that are mediated by fitness. To investigate the prevalence and characteristics of loci involved in these interactions in the budding yeast *Saccharomyces cerevisiae*, we used combinatorial DNA barcode sequencing to measure the fitness of 169 cross progeny (segregants) subjected to 8,126 CRISPRi perturbations across two environments. Global epistasis was evident in these data, with more fit segregants within each environment exhibiting greater sensitivity to genetic perturbations than less fit segregants. We dissected the genetic basis of this global epistasis by scanning the genome for loci whose effects covary with CRISPRi-induced reductions in population fitness. This approach identified 58 loci that interact with fitness, most of which exhibited larger effects in the absence of genetic perturbations. In aggregate, these loci explained the observed global epistasis in each environment and demonstrated that the loci contributing to global epistasis largely overlap with those influencing fitness in unperturbed conditions.

## Introduction

Global epistasis is a genetic background effect in which the impact of the same genetic perturbation depends on the fitness of the individual in which it occurs (Chou et al. 2011; Khan et al. 2011; Kryazhimskiy et al. 2014; Wünsche et al. 2017; Johnson et al. 2019; Rojas Echenique et al. 2019; Diaz-Colunga, Sanchez, et al. 2023; Ardell et al. 2024; Hale et al. 2024). For example, global epistasis can cause beneficial perturbations to have smaller effects in more fit individuals and larger effects in less fit individuals (Kryazhimskiy et al. 2014), while deleterious perturbations may have larger effects in more fit individuals and smaller effects in less fit individuals (Johnson et al. 2019). This phenomenon is relevant to fields such as genetics, evolutionary biology, and medicine because it influences how predictable the effects of genetic perturbations will be across individuals and populations.

Like other genetic background effects (Mullis et al. 2018; Goldstein and Ehrenreich 2021), global epistasis arises from interactions between a genetic perturbation and genetic polymorphisms, or loci, segregating within a population (Johnson et al. 2023). In the case of global epistasis, these interactions are mediated in a generic manner, in which the effects of loci on fitness, in turn, influence the impact of a perturbation (Reddy and Desai 2021; Diaz-Colunga, Skwara, et al. 2023). While researchers typically focus on how global epistasis modifies the effects of perturbations, it is also possible that the loci contributing to global epistasis have their effects altered by these interactions. In this scenario, global epistasis could shape the genetic architecture of fitness within populations, potentially altering both the effects of perturbations and the contributions of loci to fitness variability among individuals.

In populations harboring natural genetic variation, the prevalence and characteristics of loci underlying global epistasis remain empirically unresolved. Important questions include: How many loci contribute to global epistasis? What are the fitness effects of these loci in the presence and absence of the perturbations with which they interact? How do these loci collectively shape heritable phenotypic variation under perturbed and unperturbed conditions? Addressing these questions requires the ability to introduce and measure the effects of perturbations at a genomic scale in a panel of genetically diverse individuals, identify interacting loci that segregate in the panel, and assess how the effects of these loci covary with fitness.

We recently developed a chromosomally-encoded DNA barcoding system to study interactions between genetic perturbations and loci on a genomic scale in *Saccharomyces cerevisiae* (Hale et al. 2024; Levy et al. 2015; Schlecht et al. 2017). Using this system, we generated 169 barcoded segregants from a cross between the BY4716 (BY) laboratory strain and the 322134S (3S) clinical isolate (Matsui et al. 2022; Mullis et al. 2022). Into these segregants, we integrated a library of barcoded, inducible guide RNAs (gRNAs) for CRISPR interference (CRISPRi) (Qi et al. 2013; Smith et al. 2016; Smith et al. 2017). This approach produced double-barcoded strains in which the barcode identifying a segregant and the barcode identifying a gRNA were adjacent. This allowed us to perform pooled assays to measure the fitness of segregant-gRNA combinations over time via double-barcode sequencing.

Using the double-barcoded segregant-gRNA library, we demonstrated that a segregant’s fitness in the absence of perturbations (baseline fitness) is a major determinant of the effects of CRISPRi perturbations in fermentative media, the standard yeast culturing environment. Specifically, we found that deleterious perturbations had stronger effects in more fit segregants and weaker effects in less fit segregants (Hale et al. 2024). Here, we extended our CRISPRi experiment in the BYx3S cross to respiratory media, a metabolically distinct environment. We again observed a strong signature of global epistasis, with deleterious genetic perturbations showing stronger effects in more fit segregants and weaker effects in less fit segregants. These findings imply that fitness within an environment is a main determinant of how individuals respond to genetic perturbations.

In the current paper, we sought to identify the underlying genetic basis of the relationship between fitness and response to perturbation within an environment. Leveraging the continuous range of CRISPRi-induced reductions in population fitness in our pool, we developed a genome-wide strategy to map loci whose effects covary with fitness within an environment, suggesting a role in global epistasis. This approach detected 58 loci that interact with fitness across the two environments. Individually, the majority of these loci exhibited fitness effects even in the absence of perturbations and showed smaller effects when perturbations were present. Collectively, the detected loci explained the observed global epistasis.

## Materials and methods

### Generation of a double-barcoded pool of segregant-gRNA combinations

This study utilized a previously reported pool of barcoded, haploid segregants that each carry a library of barcoded gRNAs, resulting in many double-barcoded segregant-gRNA combinations (Matsui et al. 2022; Mullis et al. 2022; Hale et al. 2024). In brief, the panel of segregants was generated through the tetrad dissection of segregants from a non-barcoded cross between *fcy1Δ flo8Δ flo11Δ ura3Δ* versions of BY and 3S, both containing the same genomic landing pad for plasmid integration . These modifications provide counterselectable markers (*fcy1Δ* and *ura3Δ*) and eliminate cell-cell adhesion phenotypes (*flo8Δ* and *flo11Δ*) (Van Mulders et al. 2009).

To create the double-barcoded strains, each segregant was first transformed with a random 20-mer barcode library integrated at the landing pad, uniquely marking each genotype (Figure S1) (Gietz and Schiestl 2007). Multiple barcode integrants were retained per segregant to enable internal replication in downstream work. A second transformation was performed individually on 224 barcoded segregants to integrate a large library of barcoded gRNA plasmids, which also encoded dCas9 (Smith et al. 2016; Smith et al. 2017). This library consisted of 169 distinct genetic backgrounds including 29 segregants with at least one additional barcode. The resulting integrants from these transformations were stored in separate freezer stocks for each segregant, each containing approximately 1.25×10^8^ cells. Further details about these transformations, including selection protocols, are provided in (Hale et al. 2024).

To produce a pool of segregant-gRNA combinations, the freezer stocks of gRNA integrants for each segregant were inoculated into synthetic complete media containing glucose and lacking uracil (SC - URA). These cultures were grown to stationary phase without gRNA induction. Subsequently, 10 mL of each culture were combined together to create a ‘T0’ pool, which was immediately frozen into 250 mL ‘seed’ stocks.

### Implementation of fitness assays

Each fitness assay began by inoculating a single seed stock into 750 mL of liquid media. The cultures were grown at 30°C with shaking for 24 hours to generate the first time point (T1). Subsequently, 250 mL of the T1 culture was transferred into 750 mL of fresh media and grown for another 24 hours to produce the next time point (T2). The remaining culture was preserved as a frozen cell pellet for DNA extraction and sequencing. This process was repeated until all final time points were collected.

The same serial culturing procedure was applied to all fitness assays, with variations in the type of media (respiratory or fermentative) and the presence or absence of aTc, the CRISPRi inducer. Respiratory media consisted of SC -URA with 2% glycerol and 2% ethanol, while fermentative media used SC -URA with 2% glucose. For each environment, parallel assays were conducted with and without aTc, with harvesting occurring simultaneously. gRNA expression was induced by adding aTc at a final concentration of 250 ng/mL.

### Barcode sequencing library preparation

DNA extraction and library preparation for fitness assays in fermentative media were performed as previously described, with minor modifications to enhance efficiency and yield (Hale et al. 2024). For each time point, 120 µg of DNA was extracted from the frozen cell pellets using Zymo Research Quick-DNA Fungal/Bacterial Midiprep Kits. Double barcodes were isolated from the genome through restriction digestion with I-SceI, followed by size selection and cleanup with AMPure XP beads.

The purified double barcodes were then amplified using two PCR steps. The first PCR used Phusion polymerase (NEB), 200 ng of input DNA (53°C annealing, 60s extension, 5 cycles) and the following primers:

1. AATGATACGGCGACCACCGAGATCTACACNNXXXXNNACACTCTTTCCCTAC ACGAC
2. CAAGCAGAAGACGGCATACGAGATNNXXXXNNGTGACTGGAGTTCAGACGT GTGCTCTTCCGATCT

In the sequences above, the X’s represent Illumina multiplexing indices, and the N’s denote unique molecular identifiers (UMIs) used to eliminate PCR duplicates from the sequencing data. The product of this reaction was purified using two consecutive Zymo Research DNA Clean & Concentrator-5 columns. The purified product served as the template for the second PCR reaction, which used Phusion polymerase, 15 µL of template DNA, (53°C annealing, 60s extension, 26 cycles), and universal Illumina primers P5 and P7.

The product of the second PCR was purified using either a Zymo Research Clean & Concentrator-5 column (for fermentative media) or ethanol precipitation (for respiratory media). DNA fragments between 200 and 300 bp were extracted via agarose gel electrophoresis and recovered using Zymo Research Zymoclean Gel DNA Recovery Kits. The final product was further purified with a Clean & Concentrator-5 column, and its structure and content were verified through Sanger sequencing.

### Barcode sequencing

All time points from the fitness assays were sequenced using 150 bp paired-end reads on either a NovaSeq 6000 or HiSeq 4000 platform. A 25% PhiX spike-in was included to maintain nucleotide diversity. Custom Python scripts, run on USC’s Center for Advanced Research Computing Discovery cluster, were used to demultiplex the sequencing data and extract the barcodes. Genotype barcodes were extracted from reverse reads, while gRNA barcodes were extracted from forward reads. Reads with an average Illumina quality score below 30 across the first 35 bases of either the forward or reverse read were excluded.

Bartender, which clusters highly similar sequences and treats them as the same barcode, was then run with the merging threshold disabled (-z -1) to avoid frequency-based clustering (Zhao et al. 2018). After Bartender clustering, most reads (∼95%) perfectly matched a verified barcode sequence identified in past work (Hale et al. 2024). For the remaining cases, fuzzy matching via the process.extractOne() function in the rapidfuzz package in Python was used to associate Bartender clusters with verified barcode sequences, using a similarity score threshold of 90 or higher to define matches (Bachmann 2024). Barcode clustering and matching were performed separately for each environment (respiratory and fermentative media) and for each barcode type (segregant or gRNA).

The frequency of each double barcode across all time points and assays was calculated, excluding reads with duplicate UMIs. As in previous work, a linear model relating the frequencies of segregant and gRNA barcodes to double barcode frequencies was used to correct for chimeric amplicons (Hale et al. 2024). This model was fit using data from a separate HiSeq 4000 lane, where 10 segregants were prepared and sequenced individually, ensuring no chimera formation (Omelina et al. 2019).

### Estimation of fitness

Fitness estimation was performed as previously described (Hale et al. 2024). Frequency measurements from each time point were first normalized to account for coverage differences between sequencing lanes. For each fitness assay, PyFitSeq software was used to estimate the relative fitness of each double barcode based on its frequencies across all time points (Li et al. 2018). This software estimates the relative chance of doubling fitness of each lineage relative to the mean chance of doubling for all lineages in the whole population, based on variation in the frequencies of individual lineages. These fitness estimates are unbounded and approximately zero centered. All default PyFitSeq settings were used, other than the number of maximum iterations, which was set to 200. Because lineages represented by very few reads often result in inaccurate fitness estimates, any lineages with less than five reads at the first time point (T0) were removed from the data set, as were lineages with log likelihoods less than one interquartile range below the first quartile (Hale et al. 2024). Segregants and gRNAs with very few lineages were also removed from the data set (<100 gRNAs; <2 segregants).

PyFitSeq was performed separately for each of the four fitness assays. Within each environment (respiratory or fermentative), the resulting fitness estimates for aTc and control conditions were then normalized in order to correct out any differences between the mean fitness between assays. 83 control gRNAs that target intergenic and noncoding regions were used as a common reference point between the experimental and control assays. Six of these 83 control gRNAs were removed, as they both displayed inconsistent effects across the conditions and had fitnesses below zero indicating that they may have induced deleterious fitness effects despite not targeting a known gene. For the remaining control gRNAs, the average fitness of all lineages carrying that gRNA was determined, and the median of these averages was used to normalize the two assays.

Fitness estimation via PyFitSeq does not necessarily produce values that are perfectly centered around zero. In order to make the interpretation of fitness values more intuitive without affecting any analysis, all fitness estimates were adjusted such that the mean fitness of all lineages in the CON assay was zero. This was done for both environments individually.

### Identification and quantification of gRNA effects

The mean effects and genotype-specific effects of gRNAs in each environment were determined using mixed-effects linear models. Here, a ‘mean effect’ refers to the average effect of a gRNA across all segregants, in contrast to effects that are specific to certain segregants. Mean effect estimates for gRNAs were generated using the fixed.effects() function in nmle package in R, as well as the coef() function in R. These approaches produced identical results (Pinheiro and Bates 2000). First, the data was subset to include a specific environment (respiratory or fermentative), both of the two conditions (ATC and CON), and a single specific gRNA. The mixed-effects model *fitness ∼ segregant + gRNA + error* was fit to this subset, with the *gRNA* term indicating whether the gRNA had been induced with aTc or not. The alternate model *fitness ∼ segregant + gRNA + segregant:gRNA* + *error* was used instead if the addition of the *gRNA:segregant* interaction term significantly improved model fit. In both cases, the *gRNA* term was treated as a fixed effect and the *segregant* term was treated as a random effect. Perturbations were considered to have a potential mean effect if the p-value for the *gRNA* term was below 0.05 after Benjamini-Hochberg multiple testing correction. Perturbations were considered to have a background effect only if a significant mean effect was detected and the p-value for the *gRNA:segregant* interaction term was below 0.05 after multiple testing correction. Models were implemented using the lme() function from the nlme package in R (Gu et al. 2014). Comparison of models was performed using the anova() function in R to compare models with and without the interaction term. Additional visualizations were performed using the ggplot2() package in R (Wickham 2016).

Quantification of mean effects was performed by extracting the coefficient of the *gRNA* term from the model. A normal distribution was then generated from all gRNAs with neutral or positive mean effects. As all gRNA in our data set target genes annotated as essential in either fermentative or respiratory media, true beneficial effects are expected to be very rare, and this simulated distribution should represent the noise present in gRNA effect estimation. In order to establish a conservative threshold, a gRNA was only considered to have a mean effect in a given enironment if it both passed the p-value threshold described above and had an effect at least three standard deviations below the mean of this simulated distribution.

For gRNAs with significant background effects, model coefficients for each segregant were obtained using the predict() function in R. Segregant specific deviation from mean gRNA effects, ‘deviation values,’ for each gRNA were calculated by subtracting the overall mean effect from these segregant-specific effects. Deviation values were visualized using the superheat package in R (Barter and Yu 2017).

### Heritability estimation

Broad sense heritability for gRNA effects were computed using the replicate barcodes for both gRNAs and segregants. First control fitnesses for each segregant were obtained by averaging the fitness estimate for each segregant in the uninduced condition and control gRNAs. Then, for each gRNA that targeted an essential gene, mean segregant fitness was subtracted from each corresponding double barcode fitness in the experimental induced condition (ATC). This resulted in a gRNA effect estimate for every individual double barcode. These estimates were then used to fit the linear model *gRNA_effect ∼ genotype* + *error*. Broad-sense heritability estimates for each gRNA’s fitness effect was computed by taking the mean squares between genotypes and dividing by the total mean squares.

### Linkage mapping on deviation values

Genome-wide scans using the deviation values as phenotypes were performed for each gRNA that had a significant background effect. gRNAs with target sequences that overlap a SNP were excluded as in (Hale et al. 2024). A fixed effects linear model *(deviation ∼ locus* + *error)* was used for these scans via the lm() function in R. Significance thresholds were determined via 1,000 permutations of the data. In each of these permutations, a random gRNA was chosen and its deviation values were randomized. The lowest p-value for this randomized gRNA was saved, and the process was repeated 1,000 times. The 5th percentile of these permuted p-values was used as the significance threshold for linkage mapping. For each locus that exceeded the threshold, 2x-log_10_(p-value) drops were used to detect peak ranges, with the requirement that peaks were separated by at least 100 kb. gRNAs were removed from the analysis if their binding site was within 50 kb of a peak SNP, which affected 36 gRNAs in respiratory and 21 gRNAs in fermentative. Mapping scans and permutations were performed separately for each environment and visualized using the circlize package in R (Gu et al. 2014).

### Identification of hub loci

To identify hubs, we estimated how many loci would be expected to overlap if they were randomly distributed across the genome. This was done by dividing the genome into 20 kb bins and determining how many loci would fall within a single bin under random assortment. We then used a Poisson distribution to find the number of loci per bin that would be statistically significant. Thresholds for hubs were set at 9 and 11 for respiratory and fermentative media, respectively.

### Analysis of the relationship between locus effect and mean panel fitness

To examine the relationship between loci and fitness, we subsetted the data in each environment to lineages containing gRNAs with background effects. We then computed the least squares mean of each segregant-gRNA combination using the lsmeans() function in R. These data were split by gRNA. For each gRNA, the mean panel fitness was calculated by averaging the least squares means of all segregants carrying the gRNA. For each locus of interest, we estimated the effect of the locus on fitness in the presence of each gRNA. This locus effect was calculated as the difference between the mean fitness of segregants carrying the 3S allele and the mean fitness of those carrying the BY allele. We then computed the Spearman correlation between the locus effect and the mean panel fitness induced by a gRNA, using only the gRNAs with background effects. Spearman correlation p-values were generated using the pSpearman() function from the suppDists package in R (Wheeler and Pohlert 2024).

To examine the relationship between locus effect and mean panel fitness across the full range of mean panel fitness in our experiment, we also utilized data from additional gRNAs (equal to a quarter of the number of gRNAs with background effects) without detectable main effects or background effects. These ‘no effect’ gRNAs were selected using the sample() function in R without replacement. For these additional gRNAs, mean panel fitness and locus effect were computed as above. With this expanded dataset, we then fit linear regression models, *locus effect* ∼ *mean panel fitness* + *error*, using the lm() function in R. Estimates of locus effects for unperturbed and perturbed conditions were generated from these models using the predict() function in R. In these predictions, we set the values for mean panel fitness to zero and the 25th percentile of mean gRNA effect for gRNAs with detected background effects in a given environment.

### Genome-wide scan for loci that interact with fitness

To map loci throughout the genome whose effects interact with fitness, we extended the procedure described in the preceding section to 5,715 nuclear SNP markers genome-wide. To deal with noise at individual SNPs, we calculated mean Spearman correlations for sliding windows of 50 SNPs on the same chromosome. Significance thresholds were set within each environment using permutations in which data from gRNAs was permuted and the genome-wide scan then repeated. This process was repeated 1,000 times for each environment and the maximum absolute mean Spearman correlation was saved from each permutation. We used the maximum absolute mean Spearman correlation across all permutations as our significant threshold in each environment. Detected peaks were required to be a minimum of 50 SNPs apart, or to have mean Spearman correlations of opposite signs. We used the maximum mean Spearman correlation at each significant locus as the peak position for that locus.

### Fitness variance collectively explained by detected loci

We determined the extent to which detected loci account for observations of genetic background effects in our data. Within each environment, we first subsetted fitness data by gRNA showing a background effect. For each of these gRNAs, we then measured the extent to which all loci detected in that environment collectively explained the variance in fitness observed among segregants. To do this for each gRNA with a background effect in an environment, we constructed an additive linear model for fitness using the lm() function in R and the model *fitness* ∼ *locus_1_* + *locus_2_* + … *locus_n_* + *error*. The variance explained by each of these gRNA-specific models (R^2^) was retained and compared to the broad-sense heritability (H^2^) that had been calculated for that gRNA earlier.

### Segregant-centric and multi-locus models

For each environment, we constructed two models of fitness using the lm() function in R: a segregant-centric model and a multi-locus model. The segregant-centric model is a fixed effects model that takes into account mean panel fitness (MPF) as a continuous variable, the identity of each segregant as a categorical variable, and the interaction between mean panel fitness and each segregant: *fitness ∼ MPF + segregant + MPF:segregant* + *error*. The multi-locus model describes fitness as a function of MPF, the additive effects of all loci detected in an environment with each included as its own categorical variable, and the interaction terms for MPF and each locus: *fitness ∼ MPF + locus_1_ + locus_2_ + … + locus_n_ + MPF:locus_1_ + MPF:locus_2_ + … + MPF:locus_n_* + *error*.

Fitness estimates for each segregant under unperturbed and perturbed conditions were obtained using the predict() function in R as above. We generated 95% confidence intervals using bootstrapping, refitting each model after resampling the included gRNAs with replacement. The fitness of each segregant was then predicted with each bootstrapped model and the values retained. The 95% confidence intervals for each segregant are the 2.5th and 97.5th percentiles of predictions across 1,000 resamplings.

## Results

### Analysis of segregant-gRNA fitness in two environments

Previously, we generated a double-barcoded pool consisting of 926,939 unique combinations of gRNAs and yeast segregants, enabling the study of variation in gRNA effects across a diverse set of genetic backgrounds (Hale et al. 2024). This pool was created by integrating a barcoded plasmid library containing 8,126 inducible gRNAs targeting 1,724 distinct genes into a panel of 169 barcoded and genotyped haploid *MATɑ* BYx3S segregants (Smith et al. 2016; Smith et al. 2017; Matsui et al. 2022; Mullis et al. 2022). The chromosomally-encoded gRNA and segregant barcodes were separated by 99 bp, allowing their co-amplification and sequencing in single Illumina reads (Figure 1a and Figure S1) (Levy et al. 2015; Schlecht et al. 2017). Double-barcode sequencing of the pool at multiple time points, both with and without gRNA induction, was used to estimate the fitness of each segregant-gRNA combination (Li et al. 2018; Zhao et al. 2018). These data were further analyzed to determine the mean effect of each gRNA across segregants, the segregant-specific effects of each gRNA, and to identify gRNAs with background effects (variable effects across segregants).

**Figure 1.**
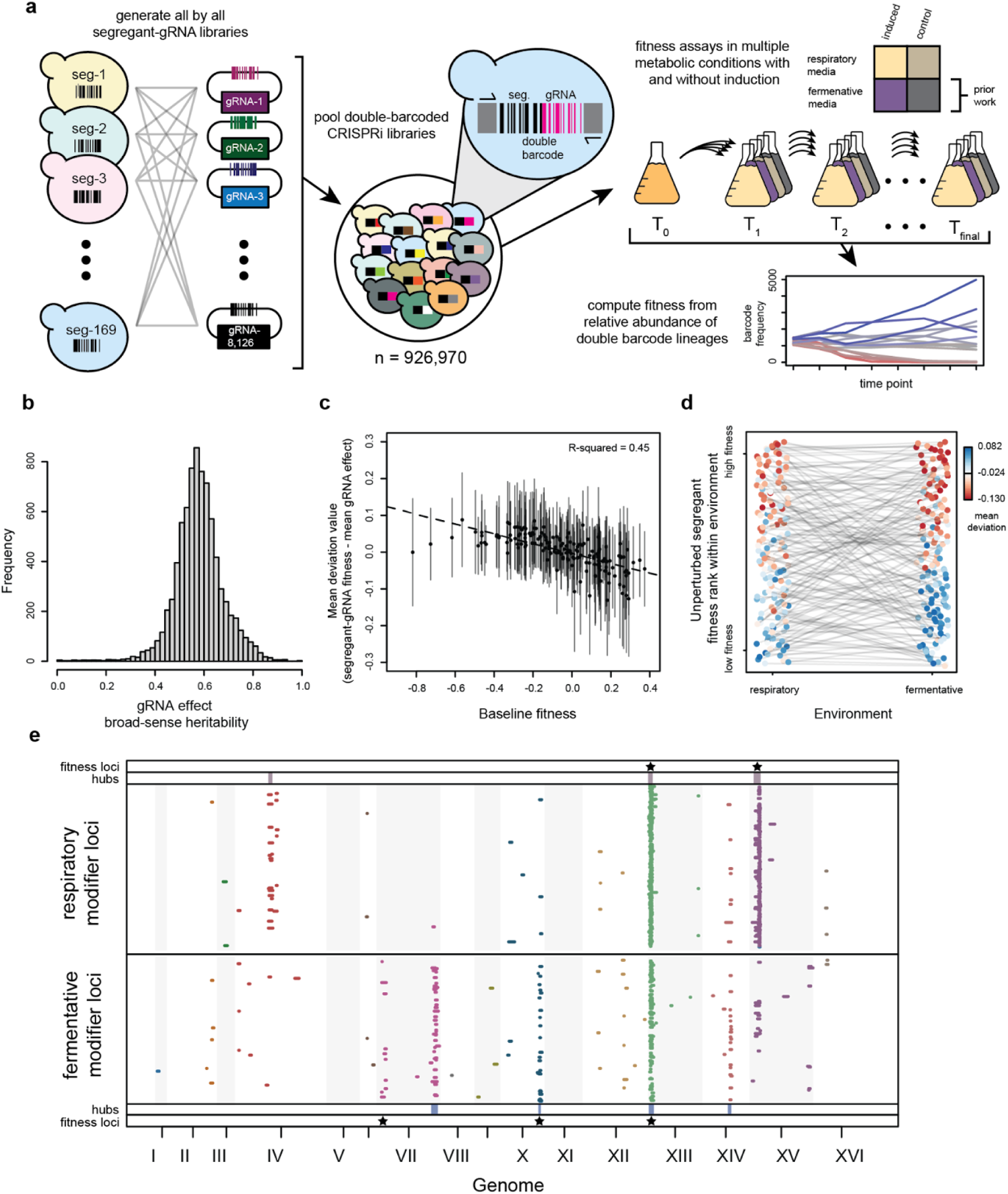
Analysis of CRISPRi perturbations in a panel of 169 yeast segregants across two environments. **a)** 926,939 unique doubled barcoded lineages, representing distinct segregant-gRNA combinations, were pooled into a single T_0_ flask. In this paper, we performed a new fitness assay in respiratory media and also reanalyzed prior data from a fitness assay in fermentative media (Hale et al. 2024). In each metabolic environment, we examined two flasks, one in which gRNAs were induced and one in which they were not. The relative fitness of each lineage within a flask was inferred from double-barcode amplicon sequencing data at multiple time points. **b)** For each gRNA exhibiting a background effect in an environment, we estimated the deviation of the gRNA’s effect in each segregant from its mean effect across all segregants. The broad-sense heritability (H²) of deviation values was calculated for all gRNAs with background effects in respiratory media (mean = 0.58, sd = 0.1). **c)** A mean deviation value for each segregant in the respiratory environment was computed by averaging all deviation values for that segregant. The relationship between baseline fitness and mean deviation values is shown (n = 169, R^2^ = 0.45, 95% bootstrap confidence interval = 0.35 - 0.57). Dots represent mean deviation values for individual segregants, with vertical lines indicating two standard deviations. This same plot for fermentative media is shown in (Hale et al. 2024). **d)** The relationship between baseline segregant fitness and mean deviation values within and between conditions is depicted. Each pair of connected dots represents a single segregant. Dot colors indicate the mean deviation value observed for a segregant in a given environment. The y-axis reflects baseline fitness rank within an environment, while the x-axis denotes the environment. **e)** Visualization of all confidence intervals for loci that show significant interactions with gRNAs, which were identified by linkage mapping. We define the confidence intervals as 2 x -log_10_(p-value) of a locus, using the p-value of the most significant SNP at a locus. Hubs for each environment are represented by bars in the rows at the top and bottom of the plot. Stars above and below mark loci detected that influence fitness in the control assays in respiratory and fermentative environments, respectively.

Fermentation and respiration represent distinct metabolic states that profoundly influence global transcription and the expression of genetic variation in *S. cerevisiae* (DeRisi et al. 1997; Rolland et al. 2002; Smith and Kruglyak 2008). In our initial study, we identified 699 gRNAs targeting 460 genes that exhibited background effects in synthetic complete medium containing glucose, a fermentative carbon source (Hale et al. 2024). In the current study, we generated fitness estimates for all segregant-gRNA combinations in respiratory media, specifically synthetic complete media containing glycerol and ethanol (Figure S2). Using these data, we identified 508 gRNAs targeting 344 genes with variable effects across segregants in respiratory media (Figure S3). These effects were highly heritable (mean broad-sense heritability [H^2^] = 0.58, sd = 0.1; Figure 1b), indicating that these variable gRNA effects have a genetic underpinning. However, because broad-sense heritability estimates depend on measurement precision, we may have failed to detect some gRNAs with variable effects that have lower heritability. Integrating the results from our previous and current studies, we identified a total of 894 gRNAs targeting 562 unique genes across both environments. Among these, 319 gRNAs targeting 242 genes exhibited background effects in both fermentative and respiratory media.

### Fitness within environment determines sensitivity to perturbation

Recent work suggests that fitness is the key factor influencing the effects of perturbations within a given environment (Ardell et al. 2024). Using data from respiratory and fermentative media, we explored this possibility and also examined the relationship between fitness and sensitivity to deleterious genetic perturbations across different metabolic environments. For all gRNAs exhibiting a background effect in an environment, we used mixed-effects linear models to estimate the deviation of a gRNA’s effect in each segregant from its mean effect across all segregants (Hale et al. 2024) (Figure S4). After calculating these ’deviation values,’ we computed a ’mean deviation value’ for each segregant in each environment by averaging all deviation values for that segregant within the given environment. These mean deviation values reflect a segregant’s relative sensitivity to perturbation compared to all other segregants.

We observed significant negative correlations between the baseline fitness and mean deviation value of segregants in both respiratory (ρ = -0.76, p-value = 3.8 x 10^-40^) and fermentative media (ρ = -0.49, p-value = 4.9 × 10^-12^; Figure 1c,d). This implies that, in both environments, perturbations had more deleterious effects in more fit segregants compared to less fit segregants. Furthermore, differences in relative segregant fitness between environments were negatively correlated with differences in mean deviation value (ρ = -0.63, p-value = 3.7×10^-20^; Figure S5). This was because segregants that were less fit in one environment and more fit in the other exhibited positive deviation values in one environment and negative deviation values in the other. In other words, the same segregant will be more sensitive to deleterious perturbations when it is placed in an environment where it is more fit and vice versa. These findings, derived from a continuous range of data in each environment, support the idea that fitness within a given environment plays a central role in determining the impact of genetic perturbations in that environment (Ardell et al. 2024).

### Genetic mapping detects a small number of loci that are linked to many gRNAs

Similar to our previous work with fermentative media (Hale et al. 2024), linkage mapping of variable responses to CRISPRi perturbations targeting the 344 genes showing background effects in respiratory media detected 397 loci in total, with approximately 1.2 interacting loci per gRNA. Based on their overlapping locations, these loci could be collapsed into three ‘hub’ loci that interacted with more gRNAs than would be expected by chance (Figure 1e and Figures S6-S7; Table 1). Our mapping resolution was insufficient to determine whether a single or multiple causative variants reside within these hubs. The newly identified hubs in respiratory media were located on Chromosomes IV, XIII, and XV and showed linkage to 18, 194 and 138 gRNAs, respectively. The hubs previously detected in fermentative media were on Chromosomes VII, X, XIII, and XIV and exhibited linkage to 39, 26, 78, and 18 gRNAs, respectively (Table S2). Two of the three hubs and in respiratory and two of the four hubs in fermentative overlap loci detected when mapping baseline fitness in the control assay (Figure 1e, Figure S8). Across gRNAs showing background effects, the mean variance in deviation values explained by the hubs was 0.14 for respiratory media (s.d. = 0.05) and 0.12 for fermentative media (s.d. = 0.04; Figures S9-S10), indicating that they account for only a small fraction of the genetic basis of the observed background effects.

### Scanning the genome for loci that interact with fitness detects more loci

Despite having nearly one million segregant–gRNA combinations in our pool, we detect only a small number of loci for each gRNA that shows a background effect. This finding suggests that the relatively small number of segregants in the pool constrained the statistical power of linkage mapping (Bloom et al. 2013). Moreover, the finding that a few hub loci interact with large numbers of gRNAs indicates that the same loci often contribute to background effects involving different gRNAs. One possible explanation for this observation is that these hubs arise due to global epistasis, with gRNAs generically interacting with hubs through their effects on fitness. In such a case, the effects of hubs might covary with the fitness perturbations of gRNAs, and additional loci undetected due to low power might also display similar patterns.

To test this hypothesis with the identified hubs, we used a new analysis framework in which we treated gRNAs as generic, quantitative perturbations of fitness, disregarding any functional information about their specific target genes (Figure 2a). For each gRNA with a background effect, we computed the mean fitness of segregants in the presence of that gRNA, which we term ‘mean panel fitness.’ This metric was continuously distributed across gRNAs in each environment (respiratory media: mean = -0.26, min = -0.66, max = -0.09; fermentative media: mean = -0.18, min = -0.69, max = -0.036; Figure S10). We then estimated each hub’s effect in the presence of each gRNA by determining the mean fitness difference between segregants carrying the 3S allele and those carrying the BY allele, referring to this measurement as the ‘locus effect.’ For each hub, this value was also continuously distributed across gRNAs in each environment (Figures S10-S11). Lastly, we examined whether mean panel fitness and locus effect were correlated across the gRNAs showing background effects. Among the seven hubs identified in the two environments, all exhibited significant Spearman correlations between mean panel fitness and locus effect (|⍴| ≥ 0.2, p-value ≤ 1.21×10^-10^; Figure 2b and Figures S12-S13).

**Figure 2.**
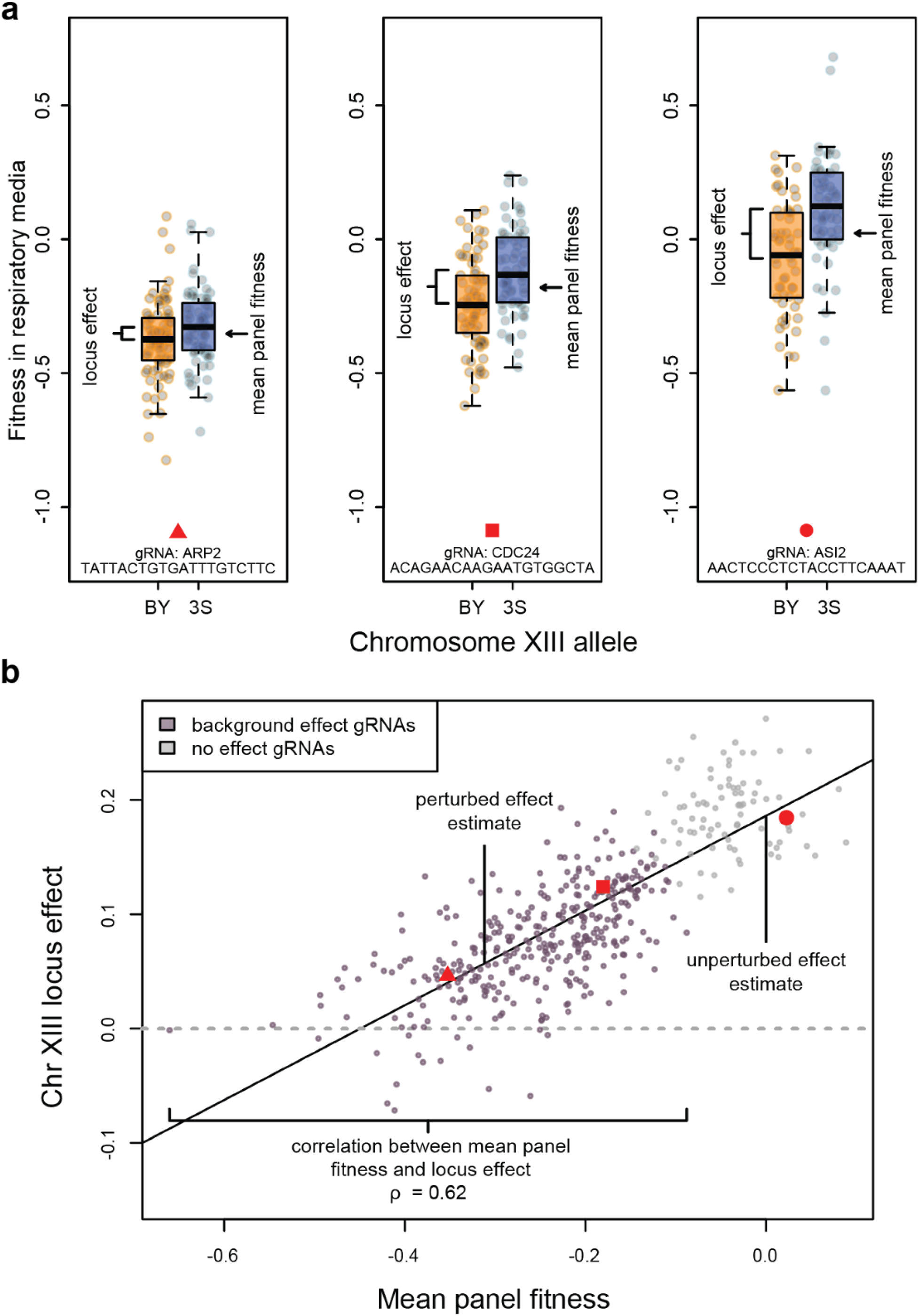
A locus whose effect covaries with mean panel fitness. In this figure, we focus on the Chromosome XIII hub in respiratory media as an example for all hubs and other loci detected by our new genetic mapping strategy. **a)** The three plots show the fitness of all segregants in the presence of three gRNAs with different effects on mean panel fitness. In each plot, segregants are grouped by their allele of the Chr XIII hub, with orange and blue indicating BY and 3S, respectively. We visualize the locus effect, which is the difference between the mean fitness of segregants carrying the 3S allele and the mean fitness of the segregants carrying the BY allele. Mean panel fitness, which is calculated as the mean fitness of all segregants in the presence of a given gRNA, is denoted with an arrow. Two gRNAs with significant background effects and one gRNA with no detected effect are shown. The boxplot central lines represent the median, while the box itself spans the interquartile range. Whiskers extend to the most extreme data points within 1.5 times the interquartile range. **b)** We show a linear relationship between locus effect and mean panel fitness. Each dot represents the locus effect measured among segregants when a particular gRNA is present. We include both gRNAs with background effects, as well as a proportional number of gRNAs that did not show significant effects. Spearman correlations were measured using only the gRNAs with background effects. However, when linear regression was used to estimate locus effects, no effect gRNAs were included to ensure the full range of mean panel fitness was represented in our models.

We next attempted to leverage the relationship between mean panel fitness and locus effect to detect new loci responsible for background effects in our study. In both environments, we scanned the genome for SNPs whose estimated effects correlated with mean panel fitness, averaging Spearman correlations within sliding windows of 50 adjacent SNPs (Figure 3a,b). We called loci significant if their absolute mean correlation exceeded the maximum absolute mean correlation observed across 1,000 permutations (respiratory threshold = 0.18; fermentative threshold = 0.15; Figure S14). At these thresholds, we detected 26 loci interacting with fitness in respiratory media and 32 loci in fermentative media (Figures S15-S21). The mean correlation at detected loci ranged from 0.29 to 0.7 in respiratory media (median = 0.46) and from 0.34 to 0.75 in fermentative media (median = 0.45). Six of the seven hubs were detected in this scan, and the remaining hub, Chr XIV in fermentative media, had a detected locus within 60 kb of the hub window, suggesting the newly detected loci are biologically relevant.

**Figure 3.**
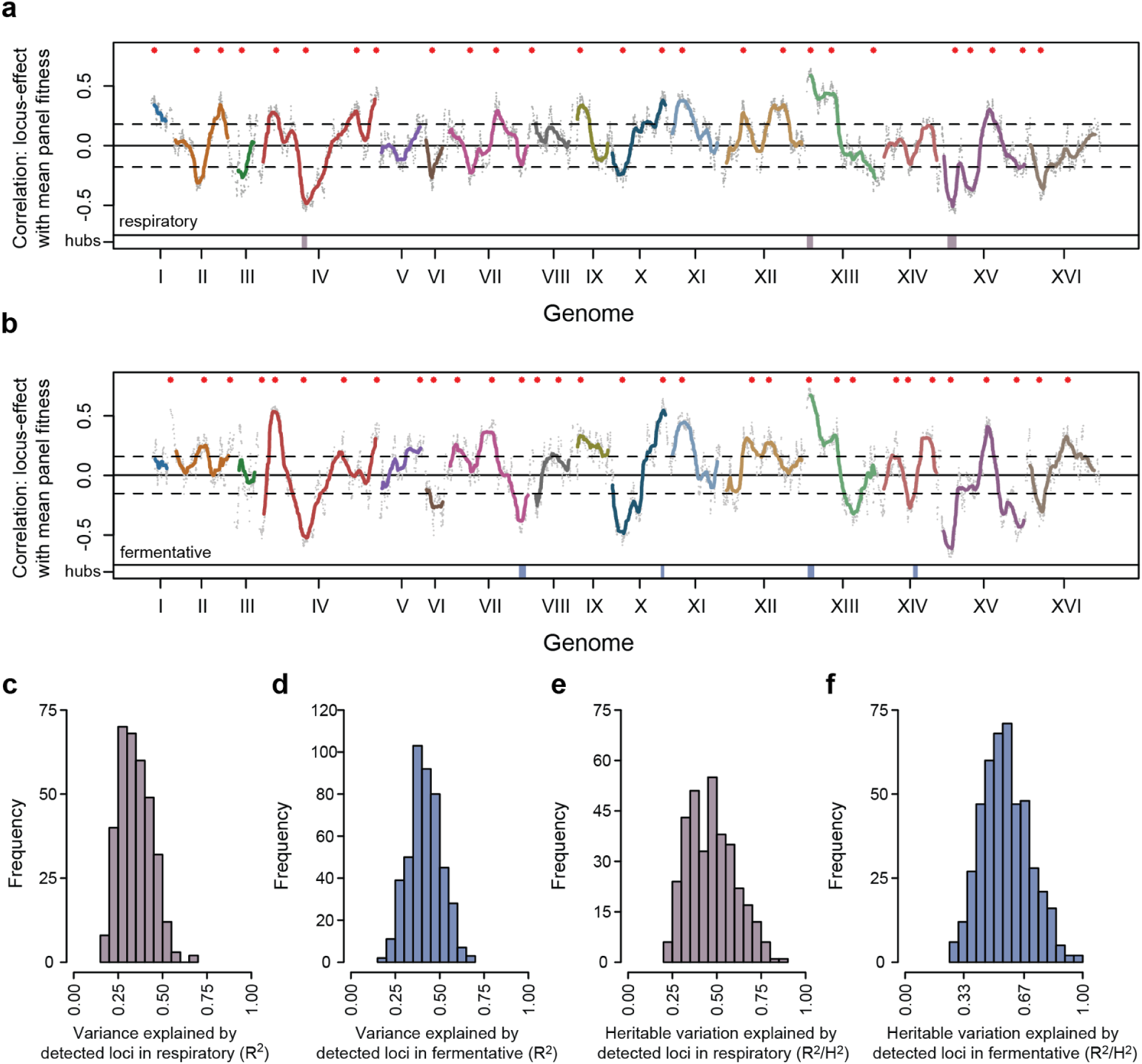
Genome-wide scans for loci that interact with fitness. **a,b)** Spearman correlations between locus effects and mean panel fitness at individual SNPs are shown as gray dots. Colored lines represent the mean correlations within 50-SNP windows. Panels (**a)** and (**b**) show results for respiratory and fermentative media, respectively. Significant loci within each environment are denoted by red dots above, with the significance threshold indicated by dashed horizontal lines. Hubs detected in each environment are indicated by bars at the bottom of each plot. **c,d)** Histograms showing the variance in fitness collectively explained by all detected loci for each gRNA exhibiting a background effect (R^2^). **e,f)** Histograms displaying the proportion of broad-sense heritability explained by all detected loci for each gRNA (R^2^/H^2^).

We assessed the extent to which the newly identified loci collectively account for the background effects in our dataset. For each gRNA exhibiting a background effect, we fit an additive fixed effects linear model of fitness among segregants as a function of all detected loci. The variance explained (R^2^) by these models ranged from 0.16 to 0.69 in respiratory media (median = 0.34) and from 0.18 to 0.67 in fermentative media (median = 0.41; Figure 3c,d). We also examined the R^2^ of these models as a proportion of H^2^ of each gRNA’s effect (R^2^/H^2^): this ratio ranged from 0.21 to 0.86 (median = 0.46) in respiratory media and from 0.26 to 1 (median = 0.55) in fermentative media (Figure 3e,f). These findings indicate that the loci identified by our mapping approach account for a meaningful portion of the observed background effects.

### Relationships between locus effects and fitness

For all detected loci in each environment, we analyzed the relationship between mean panel fitness and locus effect in greater detail using linear regression (Figure 4a,b; Figure S22). In order to examine the full range of mean panel fitness, including the absence of perturbations, we added a proportionally sized, random set of gRNAs with no detectable mean effect in an environment (referred to as ‘no effect gRNAs’). All loci produced highly significant linear regression models (maximum p-value = 1.3×10^-15^, minimum R^2^ = 0.11).

**Figure 4.**
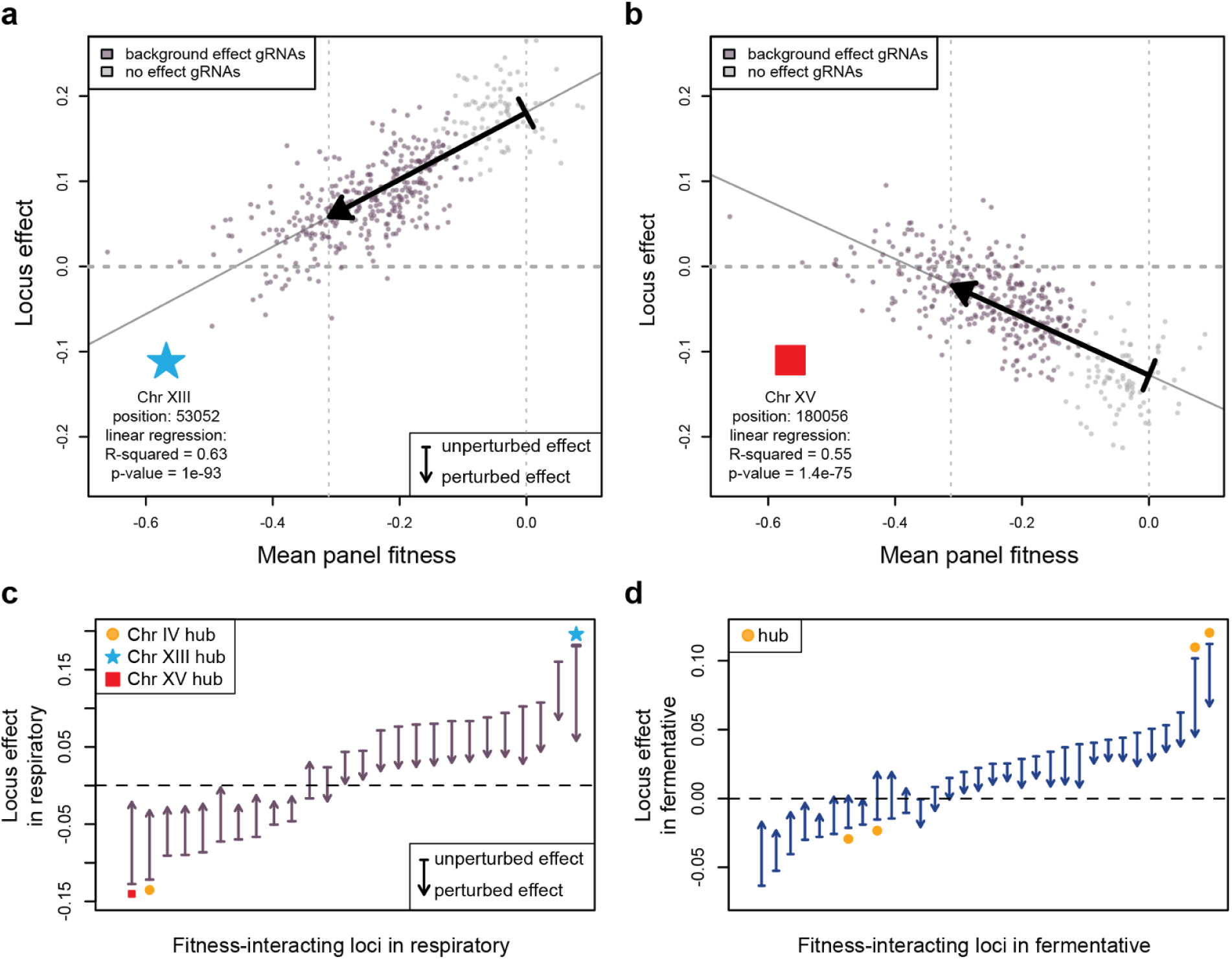
Characteristics of loci that interact with fitness. **a,b)** Visualization of linear regressions used to estimate locus effects for a given mean panel fitness. Each point represents a gRNA included in the regression with colored points indicating gRNAs with detected background effects and grey points indicating ‘no-effect’ gRNAs. Black arrow points visualize the ‘unperturbed’ locus effect estimate (defined as mean panel fitness equal to 0), while the base of each arrow represents the ‘perturbed’ locus effect estimate (defined as the 25th percentile mean panel fitness for gRNAs with background effects). The two strongest correlations in respiratory conditions are shown representing loci on Chromosomes XIII (**a**) and XV (**b**). **c,d)** The effects of detected loci in both unperturbed (arrow base) and perturbed conditions (arrow points) in respiratory (**c**) and fermentative media (**d**). Hubs are marked with symbols at the base of the arrows. The red square indicates the locus effect estimates for Chromosome XIII visualized in (**a**), the blue star indicates the locus effect estimates for Chromosome XV visualized in (**b**), and orange dots indicate all other hubs.

From the linear models, we estimated the effects of all 58 detected loci under both ‘unperturbed’ and ‘perturbed’ conditions. The unperturbed effect of each locus was defined by setting the mean panel fitness to 0, while the perturbed effect was defined as the 25th percentile of mean panel fitness among all gRNAs with a detected background effect in the respective environment (respiratory media = -0.31, fermentative media = -0.24). We selected the 25th percentile because it corresponds to a mean panel fitness value for which we have a substantial number of perturbations, but our conclusions are robust to different percentile choices.

Across the two environments, 54 of 58 loci showed larger effects in the unperturbed condition and smaller effects in the perturbed condition (Figure 4c,d and Tables S3-S4). The mean absolute effect for detected loci decreased from 0.083 to 0.027 in respiratory media and from 0.036 to 0.017 in fermentative media. Five of these loci also showed a change in sign, where the parental allele that was beneficial differed between the unperturbed and perturbed conditions. Only four loci showed a larger effect in the perturbed condition; these loci also showed changes in sign. These results show that most loci that interact with fitness in a given condition show magnitude epistasis, with smaller effects when fitness is reduced, while other forms of interaction with fitness are relatively rare (Figure S22).

### Modeling the genetic basis of global epistasis among segregants

In our study, we observe global epistasis, where each segregant responds differently to genetic perturbation in a manner related to their baseline fitness. Our mapping results suggest that these segregant-specific responses to perturbation are caused by many loci that vary in their interactions with fitness. Here, we used a series of models to evaluate whether these detected loci collectively account for the varying responses of segregants to genetic perturbation, thereby explaining the global epistasis seen in our experiment.

As a reference, we first fit a ‘segregant-centric model’ for each environment. The goal was to determine the maximum proportion of variance among segregant-gRNA combinations that could be explained by segregants’ genotypes genome-wide and the interactions of their genotypes with fitness. We applied fixed-effects linear models to describe the fitness of every segregant–gRNA combination in an environment as a function of segregant identity, the mean panel fitness resulting from a gRNA, and the interaction between these two factors (Figure 5a,b). These segregant-centric models explained 69% and 79% of the variance in fitness across segregant-gRNA combinations in respiratory and fermentative media, respectively. In contrast, simpler models that only included mean panel fitness as an independent variable respectively explained just 26% and 45% of the variance in fitness. This demonstrates that segregant identity and differential responses of segregants to perturbation account for much of the fitness variability in our experiment.

**Figure 5.**
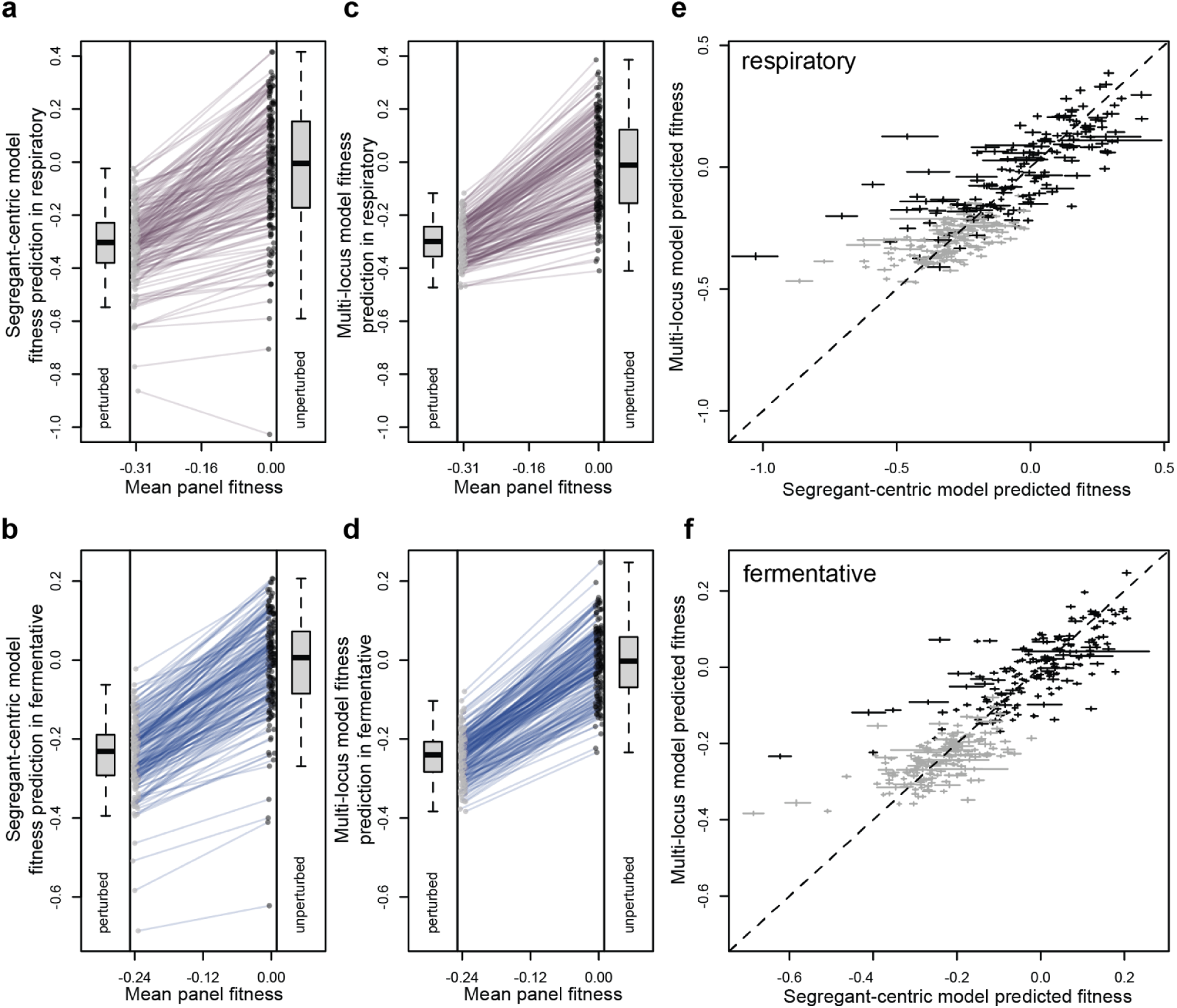
Modeling fitness and global epistasis with detected loci. **a,b)** Segregant fitness predictions for unperturbed and perturbed conditions were generated using fixed-effects linear models that describe the fitness of every segregant–gRNA combination in an environment as a function of segregant identity, the mean panel fitness resulting from a gRNA, and the interaction between these two factors (segregant-centric models). Results for respiratory and fermentative media are shown in (**a)** and (**b)**, respectively. **c,d)** Segregant fitness predictions for unperturbed and perturbed conditions were generated using fixed-effects linear models that describe the fitness of segregant-gRNA combinations as a function of a segregant’s genotype at each of the detected loci, the mean panel fitness resulting from a given gRNA, and all possible two-way interactions between loci and mean panel fitness (multi-locus models). Results for respiratory and fermentative media are presented in (**c**) and (**d**), respectively. In (**a**) through (**d**), each segregant is represented by an unperturbed value (black circles) and a perturbed value (gray circles), and the pair of values from the same segregant are connected by a blue line. **e,f)** Comparison of segregant fitness predictions for unperturbed and perturbed conditions from the segregant-centric and multi-locus models in respiratory (**e**) and fermentative (**f**) media. Horizontal and vertical lines represent 95% bootstrap confidence intervals for each segregant in the respective models. Black and gray indicate unperturbed and perturbed conditions, respectively.

We next fit ‘multi-locus models’ for each environment. These fixed-effects linear models describe the fitness of segregant-gRNA combinations as a function of a segregant’s genotype at each of the detected loci, the mean panel fitness resulting from a given gRNA, and all possible two-way interactions between loci and mean panel fitness. Multi-locus models explained 48% and 62% of the variance in fitness across segregant-gRNA combinations in respiratory and fermentative media, respectively (Figure 5c,d). While the multi-locus models explained less variance than the segregant-centric models, they still accounted for a large fraction of the observed variability in fitness. Potential explanations for the differences between these models are that additional loci that interact with fitness exist and that higher-order interactions between multiple loci and fitness play a role.

To visualize segregants’ variable responses to perturbation, we used the segregant-centric and multi-locus models to predict the fitness of all segregants in unperturbed and perturbed conditions in both environments. These predictions showed a strong signature of global epistasis, with higher-fitness segregants mostly showing higher sensitivity to perturbation than lower-fitness segregants (Figure 5a-d). Predictions from the segregant-centric and multi-locus models were highly correlated overall (ρ = 0.83, p-value = 6.0×10^-46^ in respiratory media; ρ = 0.83, p-value = 7.9×10^-44^ in fermentative media; Figure 5e,f). Correlations were high for both unperturbed (ρ = 0.79, p-value = 1.7×10^-38^ in respiratory media; ρ = 0.73, p-value = 4.7×10^-30^ in fermentative media) and perturbed predictions (ρ = 0.59, p-value = 9.6×10^-18^ in respiratory media; ρ = 0.61, p-value = 2.9×10^-19^ in fermentative media). The largest discrepancies between predictions occurred for low-fitness segregant-gRNA combinations, which are the least accurately measured in pooled assays due to biological and technical limitations. These results suggest that the identified loci play a major role in shaping fitness and global epistasis in our study.

## Discussion

We developed a genetic mapping strategy that scans the genome for loci whose effects covary with fitness, facilitating the genetic dissection of global epistasis in the context of natural genetic variation. This approach takes advantage of our experimental design in which the same panel of segregants was phenotyped across a continuous range of fitness levels induced by different CRISPRi perturbations. We identified numerous loci across the yeast genome that interact with fitness in a generic manner. Most of these loci exhibit effects in the absence of genetic perturbations and show magnitude epistasis with fitness, displaying effects that are larger at higher fitness levels and smaller at lower fitness levels. A small subset displays sign epistasis with fitness, where different alleles provide advantages under high versus low fitness conditions. Cryptic loci whose effects manifest exclusively under perturbed conditions did not appear to play a major role in our results (Rutherford and Lindquist 1998; Queitsch et al. 2002).

Interactions between loci and fitness differ from conventional gene-by-gene epistasis in which a phenotype depends on interactions between two or more specific loci (Costanzo et al. 2010; Mackay 2014; Taylor and Ehrenreich 2015; Costanzo et al. 2016; Ehrenreich 2017). However, fitness-locus interactions collectively provide insight into how global epistasis arises from conventional epistasis between genetic perturbations and many segregating loci. The identified fitness-interacting loci collectively explain why individuals’ fitness in a given environment influences their sensitivity to perturbation. These loci in aggregate make higher-fitness segregants more sensitive to deleterious perturbation than their lower-fitness counterparts within an environment. Our results demonstrate that global epistasis in the context of natural genetic variation is a complex phenomenon driven by the cumulative effects of numerous loci within an environment, each contributing to fitness and interacting with perturbations in a generic manner.

Our work ties into a broader challenge in genetics: understanding how multiple loci collectively shape complex traits (Mackay 2014; Boyle et al. 2017; Mackay and Anholt 2024). The prevailing null model for trait genetics assumes that loci contribute to traits in a predominantly additive manner (Lynch and Walsh 1998). However, the widespread presence of fitness-locus interactions raises important questions about whether similar dynamics extend to other inherited phenotypes beyond fitness. For example, in other traits, do the effects of contributing loci vary depending on an individual’s position within a trait distribution? Furthermore, how do new genetic perturbations interact with segregating loci to influence these traits? In the context of genetic diseases, our findings suggest that a single deleterious mutation with a large effect could weaken the protective impact of beneficial alleles, significantly increasing disease susceptibility. These findings emphasize the pivotal role of genetic interactions in shaping fitness, and highlight the potential broader significance of epistasis for understanding the genetic underpinnings of other traits and diseases.

## Data Availability

All raw data and code can be obtained via figshare. Processed data for respiratory media and all intermediate tables used in analyses (Supplemental Information 1-15) are provided in the supplementary information for this paper (figshare). Processed data for fermentative media are provided in (Hale et al. 2024). Raw sequencing data are available at the National Center for Biotechnology Information (NCBI) Sequence Read Archive under ascension code PRJNA1223397.

### Acknowledgments

We thank Sasha Levy, Takeshi Matsui, and Martin Mullis for their assistance in developing the segregant-gRNA pool and for past discussions related to this work, Alexandra Christensen, Matthew Dean, Michael D. Edge, Cara Hull, Zachary Krieger, Sasha Levy, Jinye Liang, Daniel Lusk, Beth Moore, David Pfennig, and Shawn Yang for providing comments on a manuscript draft, and Sergey Nuzhdin for allowing us to use his Qiagen TissueLyser.

## Funding

This work was supported by grants R35GM130381 and R56AI171091 from the National Institutes of Health to I.M.E., as well as startup funds from the University of Southern California to I.M.E.

## Author Contributions

Conceptualization: IG, JJH, IME

Investigation: IG, JJH

Formal analysis: IG, IME

Visualization: IG

Supervision: IME

Writing: IG, IME

## Conflict of Interest

None declared.

**Supplementary information is available for this paper.**

## Supplementary Materials

Figures S1 to S21

Tables S1 to S4

Supplementary Information 1 to 15

